# A human primary airway microphysiological system infected with SARS-CoV-2 distinguishes the treatment efficacy between nirmatrelvir and repurposed compounds fluvoxamine and amodiaquine

**DOI:** 10.1101/2023.06.27.546790

**Authors:** Landys Lopez Quezada, Felix Mba Medie, Elizabeth P. Gabriel, Rebeccah J. Luu, Logan D. Rubio, Thomas J. Mulhern, Jeffrey T. Borenstein, Christine R. Fisher, Ashley L. Gard

## Abstract

The COVID-19 pandemic necessitated a rapid mobilization of resources toward the development of safe and efficacious vaccines and therapeutics. Finding effective treatments to stem the wave of infected individuals needing hospitalization and reduce the risk of adverse events was paramount. For scientists and healthcare professionals addressing this challenge, the need to rapidly identify medical countermeasures became urgent, and many compounds in clinical use for other indications were repurposed for COVID-19 clinical trials after preliminary preclinical data demonstrated antiviral activity against SARS-CoV-2. Two repurposed compounds, fluvoxamine and amodiaquine, showed efficacy in reducing SARS-CoV-2 viral loads in preclinical experiments, but ultimately failed in clinical trials, highlighting the need for improved predictive preclinical tools that can be rapidly deployed for events such as pandemic emerging infectious diseases. The PREDICT96-ALI platform is a high-throughput, high-fidelity microphysiological system (MPS) that recapitulates primary human tracheobronchial tissue and supports highly robust and reproducible viral titers of SARS-CoV-2 variants Delta and Omicron. When amodiaquine and fluvoxamine were tested in PREDICT96-ALI, neither compound demonstrated an antiviral response, consistent with clinical outcomes and in contrast with prior reports assessing the efficacy of these compounds in other human cell-based *in vitro* platforms. These results highlight the unique prognostic capability of the PREDICT96-ALI proximal airway MPS to assess the potential antiviral response of lead compounds.

## Introduction

There have been over 760 million confirmed cases of COVID-19 and close to 7 million deaths worldwide (*WHO Coronavirus (COVID-19) Dashboard*, 2023). These numbers do not account for the extensive collateral damage the pandemic has caused in loss of health and livelihoods. When assessing the response from the translational science community, many positive lessons can be drawn in preparation for future pandemics such as the rapid mobilization of resources towards finding efficacious therapeutic treatment. Although COVID-19 drug discovery efforts have been met with many failures (Chakraborty et al., 2021), drug repurposing offers a cost and time saving approach. This strategy entails taking libraries of compounds already approved by governmental regulatory bodies or in the drug development pipeline for other conditions and screening them for antiviral activity. Drug repurposing can bring effective therapies to patients much earlier than the traditional 10-15 year development timeframe for new compounds, and has been demonstrated to accelerate the identification of drugs that can prevent the onset of severe symptoms or death from COVID-19 (Khataniar et al., 2022). The three FDA-emergency use authorized (EUA) drugs used in the treatment of COVID-19, remdesivir (Veklury; Gilead Sciences), molnupiravir (Lagevrio; Merck Sharp & Dohme), and nirmatrelvir + ritonavir (Paxlovid; Pfizer), were being developed for the treatment of other viral infections before being repurposed for COVID-19. Completely novel small molecules specific to SARS-CoV-2, by contrast, have not been met with great success (von Delft et al., 2023).

In this study we evaluate the antiviral activity of three compounds: fluvoxamine, amodiaquine, and nirmatrelvir (Figure 1A). Fluvoxamine and amodiaquine are FDA-approved for other indications but are exploratory antivirals for the treatment of COVID-19. They occupy two different chemical classes (Figure 1A) and, likely, two mechanisms of actions. Nirmatrelvir is the viral protease inhibitor component of Paxlovid (Joyce et al., 2022). We leveraged the 96-well format of PREDICT96-ALI (Figure 1B) and its airway-mimicking capacity to test the modulation of viral kinetics during a 6-day SARS-CoV-2 infection of the microtissues. The PREDICT96-ALI platform has previously been shown to faithfully recapitulate healthy human tracheobronchial epithelium (Gard et al., 2021) and is capable of supporting SARS-CoV-2 replication (Fisher et al., 2021, 2022). This platform has several advantages over cell culture platforms based on immortalized cell lines or other primary cell culture systems that lack key features of the host biology such as ciliary differentiation, mucociliary clearance, and the highly differentiated tissue structures and functions of the *in vivo* airway, which PREDICT96-ALI is better able to simulate. Earlier studies with PREDICT96-ALI demonstrated the ability to distinguish between antiviral efficacy of well-established therapeutics including nirmatrelvir, remdesivir and molnupiravir (Fisher et al., 2022).

**Figure 1.**
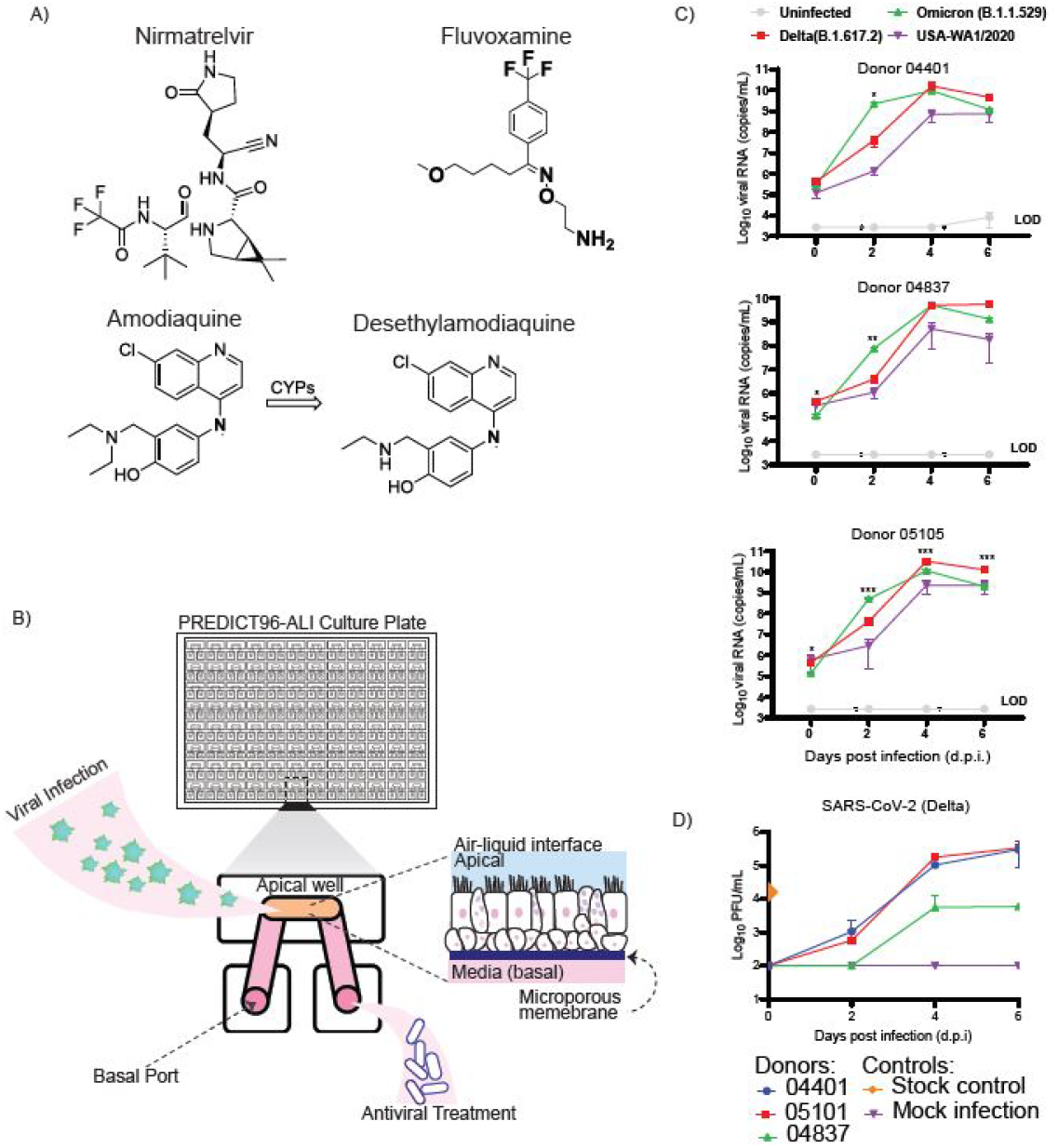
PREDICT96-ALI SARS-CoV-2 Infection and Exploratory and Control Antivirals. A) Compounds explored in this study include nirmatrelvir (an azabicyclohexane, protease inhibitor); fluvoxamine (a (trifluoromethyl)benzene, SSRI); amodiaquine (a 4-aminoquinoline, antimalarial) and its active metabolite, desethylamodiaquine. B) PREDICT96-ALI tissue culture plate containing 96 independent human bronco-tracheal model tissue atop a microporous membrane at the air-liquid interface. Tissue is infected apically for 2 hours, unbound virus is washed from the tissue, and the air-liquid interface is restored. Media and compounds are introduced to the tissue via the basal port. C) RT-qPCR analysis of apical washes of tissue derived from designated donors infected with SARS-CoV-2 variants of interest. Washes were collected from N=4 tissue replicated at 0, 2, 4, and 6 days post infection. Limit of quantification (LOQ, dotted line) indicates copy number corresponding to Ct values ≥ 37 cycles. Statistical analysis comparing SAR-CoV-2 Omicron vs Delta replication: ρ ≤ 0.05, *; ρ ≤ 0.005, **; ρ ≤ 0.0005, ***. D) Representative plaque assay for SARS-CoV-2 Delta detecting replicative virus from apical wash samples of PREDICT96-ALI airway tissue at 0, 2, 4, and 6 d.p.i. N = 3 tissue replicates per time-point and condition from one independent PREDICT96-ALI experiment.

One of the exploratory antivirals we evaluated, fluvoxamine, is an S1R agonist and serotonin re-uptake inhibitor (SSRI) used for the treatment of obsessive-compulsive disorder, depression, and anxiety disorders. Fluvoxamine can have indirect anti-inflammatory effects, which could dampen the SARS-CoV-2-induced cytokine storm (Sukhatme et al., 2021). It could also disrupt viral entry and lysosomal egress due to its lysosomotropic properties (Sukhatme et al., 2021). Preclinical data showed that in HEK293T-ACE2-TMPRSS2 cells infected with SARS-CoV-2 pseudotype virus, fluvoxamine had an IC_50_ of 10.54μM (Fred et al., 2021). Moreover, in a standard Transwell® human airway model, fluvoxamine had pronounced antiviral effects at 25μM comparable to the activity of 0.4μM nirmatrelvir. At 48hr post infection in these systems, 25μM fluvoxamine reduced SARS-CoV-2 Omicron titers by >100-fold but had highly variable (<10-fold) reduction of Delta variant viral titers (McAuley et al., 2022).

The second exploratory compound in our study is amodiaquine, a 4-aminoquinoline pro-drug and potent inhibitor of heme crystal formation in malarial digestive vacuole (Sullivan, 2017). In combination with artesunate, amodiaquine is recommended by the WHO for the treatment of uncomplicated malaria (World Health Organization, 2015). Amodiaquine shares the well-studied anti-inflammatory activity of its predecessor, chloroquine (Nagar et al., 2014) and can inhibit T-cell proliferation and IFN-γ production (Oh et al., 2016). It has also been shown to have potent antiproliferative properties against multiple cancers including colorectal carcinoma by inhibiting ribosome biogenesis (Espinoza et al., 2020) and in human melanoma cells, where amodiaquine impaired autophagic-lysosomal function through, in part, its inhibitory activity of cathepsin enzymatic activity (Qiao et al., 2013). All these properties make amodiaquine an attractive treatment option for SARS-CoV-2. Amodiaquine’s antiviral activity against native SARS-CoV-2 was first noted with a 5.15μM IC_50_ in a Vero E6 cell culture screen which tested a small subset of FDA-approved compounds (Jeon et al., 2020). A separate effort using virtual screening techniques to comb through FDA-approved compounds with structural similarity to hydroxychloroquine also predicted amodiaquine (EC_50_=0.13μM) and N-monodesethylamodiaquine to have antiviral activity (Bocci et al., 2020). This was a crucial finding since, following oral administration, amodiaquine is rapidly converted by hepatic cytochrome P450s into its most abundant metabolite, N-monodesethylamodiaquine (Li et al., 2002). Interestingly, an *in silico* drug-protein complex interaction screen searching for potential inhibitors of Mpro, the main viral cysteine protease essential for cleavage of viral polyprotein, found amodiaquine to be among the potential inhibitors (Schake et al., 2023). However, these findings regarding a potential mechanism of antiviral action directly against the virion have not been biochemically verified.

Based on the abovementioned studies, amodiaquine and fluvoxamine appeared to hold promise as efficacious antivirals and were progressed to Phase 2 and Phase 3 clinical trials, respectively. After years of work and great expenditure of resources, both compounds failed to demonstrate a reduction in viral burden in trial participants. Here, we show that when using the PREDICT96-ALI platform, these compounds had no antiviral activity, and, thus PREDICT96-ALI was able to accurately track SARS-CoV-2-infected human airway response.

## Results and Discussion

We have previously demonstrated robust native SARS-CoV-2 replication in a human primary airway microphysiological system (MPS) known as PREDICT96-ALI (Fisher et al., 2021). This result highlighted the power of PREDICT96-ALI to provide highly reproducible data while operating an MPS in a high-containment environment, a feat that no other organ-on-chip or MPS has accomplished previously. Since that time, we also reported (Fisher et al., 2022) therapeutic screening with compounds known to have moderate to high efficacy against authentic SARS-CoV-2, yielding the ability to distinguish between these compounds in a manner that tracked with clinical hospitalization data. Here, we take the process a step further by evaluating the efficacy of fluvoxamine and amodiaquine and compare it to the antiviral activity of nirmatrelvir. Fluvoxamine and amodiaquine were previously shown to have antiviral activity in other infection models, including an animal model in the case of amodiaquine, but rendered negative or inconclusive results in clinical trials.

Our previous reports of SARS-CoV-2 infection in PREDICT96-ALI (Fisher et al., 2021, 2022) employed the SARS-CoV-2 strain USA-WA1/2020. In this study, we show that airway epithelial cells derived from three donors (including a previously unreported donor, 04837) support replication of the SARS-CoV-2 Delta (B.1.617.2) and Omicron (B.1.1.529) variants up to 5-log increase in viral copy number (Figure 1C). We confirmed our RT-qPCR infection kinetic data in an additional assay by measuring the production of infectious viral particles using plaque assays (Figure 1D) which demonstrated a multi-log increase in PFU/mL over the 6-day infection time course. SARS-CoV-2 Omicron variant (B.1.1.529) showed a steeper growth curve compared to Delta (B.1.617.2) and USA-WA1/2020 in all the tissue donors tested (Figure 1C). Therefore, to capture antiviral capacity of amodiaquine and fluvoxamine, tissue infections were carried out using the SARS-CoV-2 Omicron.

Fully differentiated tissues were infected with virus at an MOI of 0.02 for two hours (h), after which unbound virus was washed off the tissue and compounds were added to the basal channel. Basal media was replaced with fresh treated media every two days. This method mirrors the likely post-viral exposure scenario experienced by infected individuals where infection occurs prior to the administration of treatment. In PREDICT96-ALI neither of the exploratory compounds, fluvoxamine and amodiaquine, showed antiviral activity over the 6-day infection and treatment period, in contrast to nirmatrelvir, which demonstrated activity even at the lowest dose tested (Figure 2A,B). This is consistent with clinical trial outcomes and contradictory to the findings reported in other comparable human lung tissue equivalent models and lung-on-chip systems (McAuley et al., 2022; Si et al., 2021), perhaps indicating that PREDICT96-ALI is able to capture nuances in tissue architecture and function and viral replication in human lung tissue.

**Figure 2.**
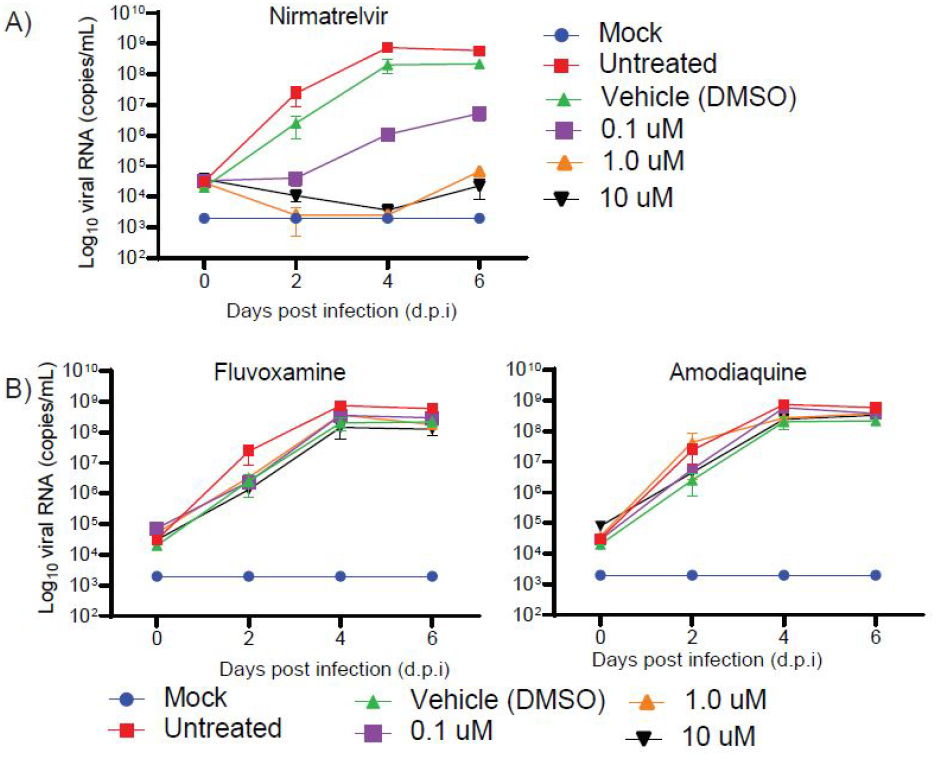
Differential response of PREDICT96-ALI to antivirals nirmatrelvir, fluvoxamine, and amodiaquine. Comparison between the antiviral effects of three drug compounds, nirmatrelvir, fluvoxamine and amodiaquine, against SARS-CoV-2 infection in primary human proximal airway tissue derived from normal human tracheobronchial epithelial cell donor 04401 grown in PREDICT96-ALI: (A) Nirmatrelvir and B) Amodiaquine and Fluvoxamine. Antiviral compounds were diluted in PneumaCult-ALI medium, added to the basal microchannel 2 h after infection and again at 2 and 4 days post-infection at 0.1, 1, or 10 µM, accompanied by mock, untreated and DMSO vehicle controls. Tissues were washed apically with HBSS to collect viral supernatant and to quantify the presence of SARS-CoV-2 viral genomes by RT-qPCR using the N1 gene target. For each condition, N=4 microtissue devices were collected. Data were collected from the same experiment and from a single PREDICT96-ALI plate and are representative of 2 independent experiments. Limit of quantification (LOQ, dotted line) indicates copy number corresponding to Ct values ≥ 37 cycles.

Outpatient studies in which individuals with confirmed SARS-CoV-2 infections self-dosed with fluvoxamine indicated no significant improvement in clinical outcomes over the standard of care. One study of 152 participants with mild COVID-19 symptoms dosed with 100mg of fluvoxamine thrice daily for 15 days resulted in clinical deterioration in 0 of 80 patients in the fluvoxamine group compared to 6 of 72 patients in the placebo group (Köhler et al., 2018; Lenze, 2021), thereby facilitating the compounds progression to larger Phase 3 trials. In one Phase 3 clinical trial, 880 participants with mild COVID-19 symptoms were given 50mg then 100mg of fluvoxamine twice daily for 15 days. In this trial, fluvoxamine reduced the level of worsening COVID-19 symptoms when compared to placebo (0.37% vs 0.73%) but treated participants had an increased risk of pneumonia; the overall risk of an adverse event was similar between the treated and placebo group (Lenze, 2022). In a larger Phase 3 trial of 1331 participants, 50mg of fluvoxamine was given twice daily for 10 days; 26/674 participants (3.9%) in the fluvoxamine group were hospitalized, had an urgent care visit, had an emergency department visit, or died compared with 23/614 participants (3.8%) in the placebo group (McCarthy et al., 2023; Naggie, 2023). The authors of the latter study did not recommend fluvoxamine at the dose and duration tested for patients with mild to moderate COVID-19.

With amodiaquine, we were also unable to detect any significant reduction in viral loads even at the 10μM highest dose tested (Figure 2B). This contrasts with a previous report from another institution using an alternate lung-on-chip model infected with SARS-CoV-2 pseudovirus, where 1.24μM amodiaquine reduced viral entry by 59.1% and 1 μM desethylamodiaquine reduced entry by approximately 60% (Si et al., 2021). The authors went on to show that amodiaquine, when administered prophylactically or post-infection, reduced viral load by ∼70% in the lung homogenates of hamster infected with native SARS-CoV-2 and noted a downregulation of genes associated with an inflammatory response. A modest Phase 2 clinical trial consisting of 186 symptomatic participants commenced where 39 participants received the drug regimen artesunate-amodiaquine at 200/540 mg once daily for 3 days (Si et al., 2021). According to the authors, at this concentration the estimated levels of desethylamodiaquine in the lung should exceed 50% the estimated inhibitory concentration for SARS-CoV-2. Unfortunately, no significant improvement in viral clearance was found when compared to standard of care (1000mg of paracetamol 6-hourly as needed). Due to the sample size, the authors did not draw any definitive conclusions (Chandiwana et al., 2022) and a larger follow-up study was recommended. Nevertheless, evidence is emerging of amodiaquine’s ability to inhibit human cathepsin B which could decrease the infectivity of several category A and B pathogens that enter host cells via endosomes (Zilbermintz et al., 2015). Despite inconclusive COVID-19 clinical findings, amodiaquine may yet show promise as an anti-infective for other pathogens.

Although prospects for success are limited and the process daunting, especially in the time-sensitive context of a pandemic, drug repurposing has many potential advantages. Potentially efficacious treatments that have already undergone rigorous process and manufacturing development as well as toxicity and tolerability studies may make their way to patients much faster, reserving valuable time and resources to *de novo* drug development. A primary hurdle to efficient drug repurposing in the challenging context of an emerging pandemic is a lack of high-fidelity, efficient and predictive preclinical drug development tools or reliable animal models (Sultana et al., 2020). Here, we evaluate three repurposed compounds that have received extensive clinical evaluation, one with known efficacy and two with inconclusive or absent levels of efficacy, in our novel high-throughput, high-fidelity PREDICT96-ALI MPS system. Previous reports with existing *in vitro* or MPS models incorrectly predicted efficacy in the clinically ineffective compounds, highlighting a gap in the availability of predictive preclinical tools for the COVID-19 pandemic. The results of our studies using PREDICT96-ALI are consistent with the results of clinical trials for the repurposed compounds, a capability that leads to rapid advancement of successful treatments and avoidance of wasted resources for those that ultimately prove ineffective. Features of PREDICT96-ALI that allow it to resolve drug compound efficacy and account for the observations of this study may be due to a combination of two factors. First, PREDICT96-ALI features human tissue derived from primary airway epithelial cells and matures the tissue with dynamic fluid shear stress control, permitting homogeneous nutrient distribution, elimination of chemical gradients and sufficient oxygenation of the culture media. Second, use of native SARS-CoV-2 in a high-containment laboratory enables evaluation of the full viral life cycle, which is critical to assessing compounds with varied mechanisms of action. This ability to accurately predict clinical outcomes using an MPS represents a new and major milestone in the effort to establish human primary cell-based preclinical models as a tool to address emerging infectious disease threats and more broadly in the drug development pipeline.

## Methods and Materials

### Microfluidic Platform and Integrated Micropumps

The PREDICT96 microphysiological system consists of a microfluidic culture plate with 96 individual bilayer devices (Figure 1B) and a recirculating perfusion system driven by 192 pneumatically-actuated micropumps embedded in the plate lid (Azizgolshani et al., 2021). Because the apical well chamber is at an ALI, only 96 pumps are utilized in the PREDICT96-ALI platform where the micropumps transfer media between the inlet and outlet wells corresponding to the ports of the basal microfluidic channel, establishing a hydrostatic pressure differential between ports, which results in media flow through each microchannel during tissue maturation.

### Preparation of Human Primary Tracheobronchial Epithelial Cells from Healthy Living Donors

The primary normal human tracheobronchial epithelial cells (NHBE) used included Donor 05101, a 29 year old Caucasian female (non-smoker), Donor 04282, a 23 year old Caucasian male (marijuana smoker), and Donor 04401, a 16 year old Caucasian male. All donors were sourced from Lifeline Cell Technology. NHBEs were seeded in PREDICT96-ALI plates as previously described (Gard et al., 2021). Briefly, each device was seeded directly on the PREDICT96-ALI membrane in the apical chamber with 10,000 cells in Small Airway Epithelial Growth Medium (Lonza) containing 100 U/mL penicillin–streptomycin (Thermo Fisher), 5 μM ROCKi (Tocris), 1 μM A-83-01 (Tocris), 0.2 μM DMH-1 (Tocris), 0.5 μM CHIR99021 (Tocris) (hereafter referred to as SAGM + 4i). Tissues were grown for 48 h in SAGM + 4i media followed by 48 h of submerged tissue differentiation in HBTEC-ALI media (Lifeline Cell Technology) including 100 U/mL penicillin/streptomycin. Tissues were placed at an air-liquid interface (ALI) by aspirating all media from the apical well chamber of each device and replacing the media in the basal channel and wells with PneumaCult-ALI (STEMCELLTechnologies) and were cultured at an ALI at 37°C in 5% CO_2_ and 90% humidity for 3-4 weeks prior to infection with fresh media changes every two days.

### Inoculation of PREDICT96-ALI Tissues with SARS-CoV-2

All SARS-CoV-2 infections were performed in a Biosafety Level 3 (BSL3) facility at the New England Regional Biosafety Laboratory at Tufts University with appropriate institutional biosafety committee approvals. SARS-CoV-2 isolate USA-WA1/2020 passage 4 virus was obtained from Joseph Brewoo and Sam R. Telford III (New England Regional Biosafety Laboratory, Global Heath & Infectious Disease, Tufts University). Working stocks of USA-WA1/2020 strain was generated by propagating the virus on Vero E6 cells (ATCC, USA) as previously described (Fisher et al., 2021). Delta variant lineage B.1.617.2 (source: hCoV-19/USA/MD-HP05647/2021; BEI Resources, USA) and Omicron variant lineage B.1.1.529 (source: hCoV-19/USA/GA-EHC-2811C/2021; BEI Resources, USA) were passaged in Calu-3 cells as recommended by BEI Resources. Inoculation of PREDICT96-ALI proximal airway tissues with SARS-CoV-2 were performed using SARS-CoV-2 (Figure 1B) diluted to the desired 0.02 MOI. On 2, 4, and 6 days post infection (d.p.i.), the apical surface of the tissues was washed twice with HBSS to harvest virus. Fresh PneumaCult-ALI media was introduced into the basal microchannel before returning the plate to 34°C at 2 and 4 d.p.i.

### Antiviral dosing

Nirmatrelvir (MedChemExpress), Fluvoxamine maleate (Selleck Chemicals), and Amodiaquine dihydrochloride dihydrate (Millipore-Sigma) were diluted fresh from 10 mM stock concentrations in dimethyl sulfoxide (DMSO) to 10, 1 or 0.1 µM in complete PneumaCult-ALI. DMSO diluted in media served as the vehicle control. Drugs and controls were applied in the basal microchannels (Figure 1B) of PREDICT-ALI proximal airway tissues 2 h after infection and at 2 and 4 d.p.i. All drug dosing conditions were performed in quadruplicate.

### RT-qPCR

HBSS washes from PREDICT96-ALI devices were tested by reverse transcription quantitative polymerase chain reaction (RT-qPCR). Briefly, direct RT-qPCRs were performed on heat-inactivated samples without extraction using the QuantiTect Reverse Transcription Kit (Qiagen). The reactions were run in an Applied Biosystems QuantStudio 7 Flex System (Thermo Fisher Scientific) using the following primers and probes targeting SARS-CoV-2 nucleocapsid protein: nCOV_N1 Forward Primer Aliquot, nCOV_N1 Reverse Primer Aliquot, and nCOV_N1 Probe Aliquot (IDT). Samples with cycle threshold (ct) values above 37 were omitted.

### Plaque assay

Hanks’ Balanced Salt solution wash samples collected from inoculated PREDICT96-ALI plates were titered via plaque assay as previously reported (Jureka et al., 2020). Briefly, Vero E6 cells (ATCC, USA) were plated in 6-well plates and incubated at 37°C and 5% CO_2_ until confluent. HBSS wash samples were initially diluted 1:5 or 1:10, and Vero E6 cells were incubated with viral dilutions for 2 h then washed with PBS, overlaid with DMEM and 1.2% Cellulose (colloidal, microcrystalline (Sigma-Aldrich)), and incubated for 3 days. Plaque forming units/milliliter (PFU/mL) were calculated as follows: number of plaques / (dilution x total inoculum volume (mL)).

### Statistical analysis

Statistical analysis comparing SAR-CoV-2 Omicron vs Delta replication was performed in GraphPad Prism software version 9.3.0 (Graphpad Software, Inc.). RT-qPCR data were log-transformed and analyzed using a two-way analysis of variance (ANOVA) with Dunn–Šidák’s test for multiple comparisons (noted in figure legend). *A*ll ρ-values signified as follows: ρ ≤ 0.05, *; ρ ≤ 0.005, **; ρ ≤ 0.0005, ***.

## Acknowledgments

The authors gratefully acknowledge Roger Odegard, Timothy Petrie, and Alla Gimbel for overall technical and management support, John Tonkiss for biosafety support, Rick Crocker and Karen Arenburg for support of the laboratory infrastructure, Robert Gaibler, Yazmin Obi, and Jonathan Crocker for laboratory support, and Else Vedula for key technical guidance and critical review of the manuscript. We gratefully acknowledge the technical and programmatic support of Sam Telford, Joseph Brewoo, Donald Girouard, Wanda Girouard and the staff at the Tufts University New England Regional Biosafety Laboratory.

